# Susceptibility of ecosystems to interaction timing

**DOI:** 10.64898/2026.04.06.716858

**Authors:** Phillip P.A. Staniczenko, Jilles Verwoerd, Berry J. Brosi, Debabrata Panja

## Abstract

The phenology of organisms worldwide is shifting in response to changes in environmental conditions. There is growing concern that resulting timing mismatches among interacting species will negatively impact system-level properties, yet there is no general framework for evaluating community responses to changes in phenology. To address this gap, we developed a mathematical framework based on local stability analysis and used it to assess the resilience implications of phenological perturbations with a multi-year, highly time-resolved empirical dataset on subalpine plant-pollinator communities. The forecasted effects of phenological perturbations were largely independent of perturbations to species densities, indicating the potential for even small changes in phenology to disrupt the functioning of ecosystems that are otherwise highly stable.

---

Phenology, the timing of biological events, is changing globally (Parmesan 2007, Thackeray et al. 2016, Roslin et al. 2021). In ecological communities, species respond to different abiotic cues that are becoming increasingly decoupled from one another (Bartomeus et al. 2011; Ovaskainen et al. 2013). As phenology determines when species are active, changes to longstanding temporal patterns can cause timing mismatches that impact the chance and frequency of functionally-important interactions (Diez et al. 2012; Theobald et al. 2017; Carter et al. 2018; Duchenne et al. 2020). Plants flowering earlier than usual, for example, can reduce temporal overlap with their insect pollinators, lowering effective reproduction rates in plants and resource availability to pollinators, thereby causing decreases in population numbers in subsequent seasons (Ashman et al. 2004; Kudo et al. 2004; Memmott et al. 2007; Rafferty & Ives 2011; Kudo & Ida 2013).

Although the effects of phenological shifts on pairwise and one-to-many species interactions are increasingly being studied, the effects on whole-community dynamics are less understood. A promising starting point is local stability analysis (May 1972; Allesina & Tang 2015) because phenology is known to shape the structure of ecological networks (Peralta et al. 2020; Peralta et al. 2024) and local stability analysis connects the structure of ecological networks to the capacity for communities to recover from externally-driven perturbations. Early work assumed random network topologies and random combinations of interaction types (e.g., trophic, competitive, mutualistic) (McCann 2000), while more recent studies have investigated the stabilizing (or destabilizing) roles of interaction type (Allesina & Tang 2012; Stone 2020), strength (Brose et al. 2006; Gellner & McCann 2016), and topology (Jacquet et al. 2016; Anderson et al. 2024). This additional realism has helped reveal many of the underlying factors influencing ecosystem stability, but there have been two major impediments to examining the role of phenology: (i) existing approaches do not account for seasonality, instead assuming that species are always active and available to interact during a season, and (ii) only the effects of perturbations directly affecting species densities have been considered (Ramos-Jiliberto et al. 2018; Glaum et al. 2021; Duchenne et al. 2022).

Recent advances in the collection of highly time-resolved network data have made it possible to investigate the system-level effects of phenological shifts. Here, we developed a local stability analysis framework that considers perturbations to species phenologies in addition to perturbations to species densities. The framework also incorporates a recently proposed inverse problem method (Gellner et al. 2023) to infer hard-to-measure empirical interaction strengths from more readily collectable field data on relative species abundances. We used a uniquely detailed eight-year, three-site data set on plant-pollinator interactions (Morozumi et al. 2022) to assess the correlation between an ecosystem’s response to phenological perturbations and its response to perturbations to species densities.

There are three possible system-level outcomes. The most expected outcome is that ecosystem responses are aligned such that the more stable an ecosystem is to perturbations to species densities, the less susceptible it is to phenological perturbations (H1). This follows the reasoning that resilient systems, so far measured primarily by their ability to recover quickly from the loss of individuals, have inherent features that make them able to withstand different kinds of perturbations. Alternatively, the less expected outcome is that responses are anti-correlated—that is, there are system-level trade-offs such that increased resilience to perturbations to species densities comes at the cost of increased susceptibly to phenological perturbations (H1A). The third option is that there is no appreciable relationship between the two responses (H0). In addition to system-level relationships, we also aimed to correlate species-level responses with organismal and network properties, with the expectation that species with fewer interaction partners, i.e., specialists, would be more sensitive to shifts in phenology than generalists (Wilson & Yoshimura 1994, Miller-Rushing et al. 2010, Brosi 2016, Peng et al. 2025).

## Population dynamic model

We described underlying coupled plant and pollinator dynamics using a modified version of the consumer-resource model by Holland & DeAngelis (2010). The model assumes that plant and pollinator species both benefit from an interaction (facultative mutualism), but that plants also incur a cost to producing floral resources and nectar to attract insects. We modified the base model to include a new parameter, *ϕ*, that accounts for the timing constraints on interactions imposed by each species’ individual phenology:

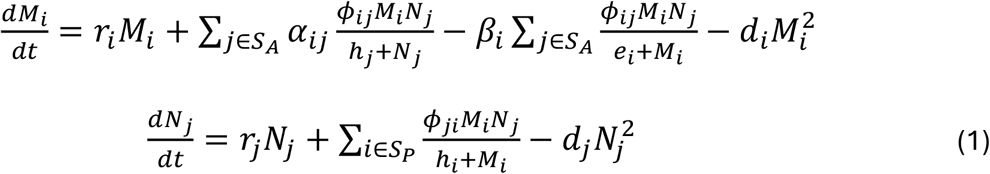

where *M*_*i*_ and *N*_*j*_ represent the densities of plant (*i* ∈ *S*_*P*_) and pollinator species (*j* ∈ *S*_*A*_), respectively, with intrinsic growth rates given by *r*_*i*_ and *r*_*j*_, and death rates by *d*_*i*_ and *d*_j_. The effect of interactions on population growth is represented by *α*, with non-zero and positive values for *α*_*ij*_ if plant species *i* and pollinator species *j* are involved in a mutualistic interaction; *h*_*i*_ and *h*_*j*_ are half-saturation constants. For plants, the cost to providing resources to pollinators is represented by *β*_*i*_; *e*_*i*_ is a half-saturation constant. The phenology parameter, *ϕ*, rescales the expected number of encounters among interacting species pairs according to the amount of temporal overlap between the two species:

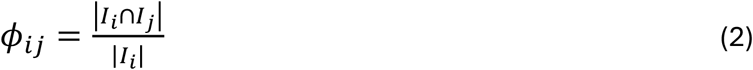

where *I*_*i*_ is the length of time that focal species *i* is active and *I*_*j*_ is the length of time that its interaction partner is active; values for *ϕ*_*ij*_ range from 1 if *j* is active the entire time *i* is active, to smaller values the shorter the fraction of time that *j* overlaps with *i*. The phenology parameter is not symmetric, i.e., *ϕ*_*ij*_ ≠ *ϕ*_*ji*_, and has a minimum value of 0 if two species, even if known to interact in other contexts, are not active concurrently in a season at a given location. By contrast, modelling approaches that do not account for phenology implicitly assume *ϕ*_*ij*_ = 1 for all interacting pairs of species, i.e., all interaction partners are always present and available to interact. Note that *ϕ*_*ij*_ does not rescale mutualistic interaction strengths (*α*_*ij*_), nor the cost of attracting pollinators (*β*_*i*_), but simply the expected number of interactions between two species, i.e., *ϕ*_*ij*_ acts on all terms involving the product *M*_*i*_*N*_*j*_.

## Local stability analysis

Local stability analysis traditionally follows a forward modelling approach in which interaction strengths, *α*, and species-specific parameters (*r*_*i*_, *d*_*i*_, etc.) are treated as fixed inputs. These parameters are used to solve for a system’s steady-state densities, with stability assessed by evaluating whether (and, if so, how quickly) the system will return to its equilibrium state following an infinitesimally small perturbation to steady-state densities (May 1973). Measuring interaction strengths in the field is challenging, so studies have typically sampled *α* values from plausible synthetic distributions rather than using empirically-informed values (Allesina & Tang 2015). Recently, however, inverse problem methods have been proposed for inferring interaction strengths from species abundance data, such that *α* is no longer treated as a fixed input (Gellner et al. 2023). With an inverse problem method, steady-state densities are estimated from empirical data and then combined with the traditional set of species-specific parameters to constrain a population dynamic model (e.g., Eqn 1) and solve for *α*. Solving such an inverse problem typically means solving an underdetermined set of equations (de Ruiter et al. 1995; Neutel et al. 2007), resulting in many plausible sets of interaction strengths that satisfy the equilibrium condition (see Methods).

For a given set of inferred interaction strengths, we assessed conventional local stability by performing an eigenvalue decomposition of the community matrix (May 1973), the Jacobian matrix of a population dynamic model evaluated at equilibrium, i.e., at steady-state species densities. In this case, partial derivatives in the Jacobian matrix are taken with respect to changes in species densities, and stability is determined by the real part of the dominant (largest) eigenvalue: if the value is negative, then the system is stable and will eventually return to its equilibrium state following a perturbation; if the value is positive, then the system is unstable and will transition to a different state with potentially different community composition. To assess the susceptibility of a dynamical system to changes in phenology, we considered a complementary Jacobian matrix in which partial derivatives are calculated with respect to changes in *ϕ* rather than changes in species densities (see Methods). This Jacobian matrix evaluated at equilibrium species densities, which we refer to as the B matrix, describes individual species sensitivities to phenology. Each element quantifies how the growth rate of a particular species is expected to change *immediately following* an infinitesimally small decrease in its temporal overlap with one of its interaction partners; in this regard, the B matrix describes responses that occur on short timescales, as with reactivity (e.g., Tang & Allesina 2014). Here, we focused on stability because reactivity does not directly inform whether a system will return to its original equilibrium point following a perturbation to species densities. The elements of the B matrix can be positive, signifying initial increases in focal species densities, or negative, signifying initial decreases in densities.

As the B matrix is rectangular (species-by-interaction) rather than square (as with the species-by-species community matrix), eigenvalues cannot be calculated for the B matrix. Instead, we performed singular value decompositions (Golub & Van Loan 1996) and used the dominant (largest) singular value as a measure of system-level susceptibility to changes in phenology. Note that while the elements of the B matrix can be positive or negative, singular values of the B matrix are necessarily positive due to the matrix multiplication steps used to calculate them. Here, the dominant singular value describes the largest collective first-order, initial response of species to phenological changes. Because singular values are always non-negative real numbers, there is no equivalent transition from stable to unstable as there is with the dominant eigenvalue crossing from negative to positive. This is why we use “susceptibility” when referring to phenology-based ecosystem responses, as in the *potential* for large changes in species densities following perturbations to the timing of interactions.

## Mathematical expectation

We found that, from a mathematical perspective, there is no *a priori* reason to expect a strong relationship between dominant eigenvalues and dominant singular values, i.e., mathematical expectations are consistent with H0. However, this does not rule out the possibility for strong positive (H1) or negative (H1A) correlations between dominant eigenvalues and dominant singular values in empirical data, especially as both types of values are determined by the same set of mechanistic parameters in Eqn 1. The full mathematical argument is given in Supplementary Materials, with the central idea that while the community matrix focuses on interspecific feedbacks mediated through interactions, the B matrix instead summarizes isolated, species-specific responses to changes in temporal overlap with interaction partners. As such, although both lines of inquiry sit within a local stability analysis framework and involve similar mathematical and computational steps, they measure fundamentally different response pathways to externally-driven perturbations.

## Ecosystem susceptibility to phenological shifts

We used replicate plant-pollinator network data, collected weekly throughout the growing season for eight years (2016–2023) at three field sites (“AP,” “GT,” and “VB”) in and around the Rocky Mountain Biological Laboratory, Colorado, USA, to parameterize the population dynamic model (Eqn 1) and infer interaction strengths. For each site, the empirical data specified binary network structure, phenological coefficients (*ϕ*), and estimates of inter-season equilibrium species densities; all other species-specific parameters (e.g., intrinsic growth rates) were set to constant values. We assumed each site was at steady state and stable (by tuning death rates, see Methods), and inferred interaction strengths using hit-and-run sampling. For each set of sampled *α* values, we computed the dominant eigenvalue from the corresponding community matrix and the dominant singular value from the corresponding B matrix (see Methods).

We used the distributions of dominant eigenvalues (for stability) and dominant singular values (for susceptibility) generated from multiple sets of sampled *α* values for each site to evaluate ecosystem resilience to externally-driven perturbations. Our implementation of the inverse problem method only stipulated that dominant eigenvalues have negative real part, so the resulting distributions of dominant eigenvalues (and therefore corresponding distributions of dominant singular values) reflect natural variations for each site. This site-specific approach is preferable to cross-site analyses due to the large number of confounding factors that are known to influence stability. The different levels of stability considered for each site corresponded to the different configurations of interaction strengths consistent with steady-state dynamics, with the inverse problem method providing a built-in way of controlling for empirical network topology and constraints on the positioning of different interaction types.

There was weak positive correlation between dominant eigenvalues and dominant singular values for each of the three sites (AP in Fig. 1, plots for GT and VB sites in Supplementary Materials; Spearman’s rank correlation for GT: *r* = 0.297; and for VB: *r* = 0.008; *n* = 100,000 samples for each site). This result lends support to H0 and followed the mathematical expectation. The correlation for the GT site suggested slight support for H1 (more stable systems are less susceptible to phenological perturbations); results did not support H1A (trade-off between stability and susceptibility). Dominant eigenvalues describe the exponential return time to equilibrium following a perturbation to species densities, while estimated responses to phenological perturbations scale linearly with dominant singular values. Across field sites, dominant eigenvalues represented a similar range of stability levels and singular values spanned a two-to three-fold range of effect sizes.

**Fig. 1.**
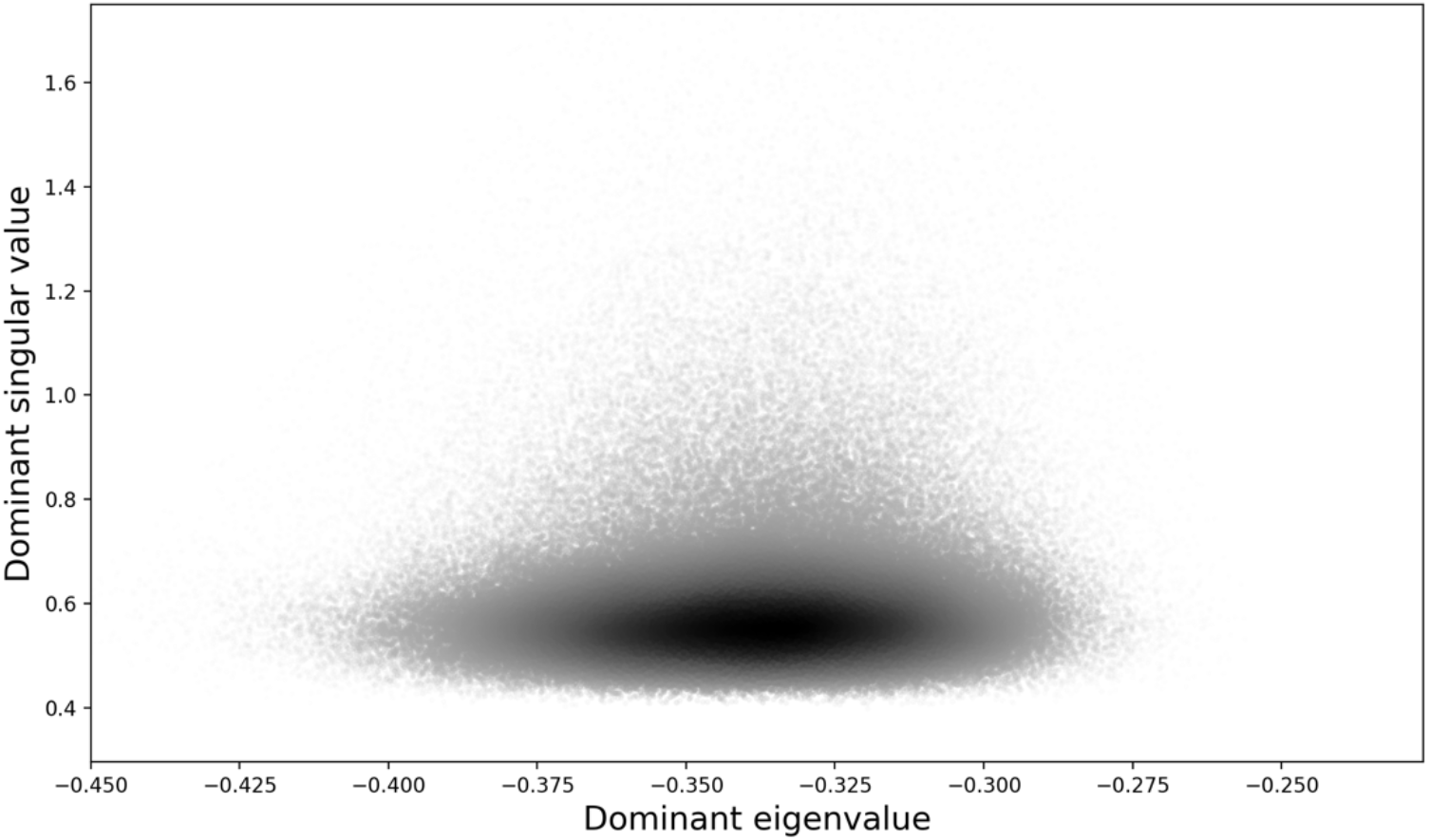
System-level stability assessment. The correlation between dominant singular values of the B matrix and dominant eigenvalues of the community matrix is extremely weak for the AP site (Spearman’s rank correlation: *r* = 0.051). Each point in the scatter plot represents a dominant singular value and dominant eigenvalue pair corresponding to a distinct set of inferred interaction strengths (*n* = 100,000 samples); darker areas indicate greater densities of points.

## Species-level sensitivity

We obtained phenological sensitivity values for each species for each field site in two steps: (i) we averaged over multiple inferred sets of *α* to obtain a representative species-by-interaction B matrix and then (ii) for each species (row) in the B matrix, we took the arithmetic mean across all observed interactions (columns) and divided by the species’ density to obtain per capita sensitivity values. The sign of a sensitivity value follows from the dynamical model, Eqn 1. Since plants are modeled as experiencing both costs and benefits to an interaction, decreases in temporal overlap with interaction partners can result in either an initial increase (positive sensitivity values) or decrease (negative sensitivity values) in population densities. Pollinators only receive benefits from an interaction, so decreases in temporal overlap always result in an initial decrease in population densities, with larger initial decreases in population densities associated with more-negative sensitivity values.

To understand what drove variation in sensitivity values for the three sites, we examined the influence of two functionally-relevant organismal and network properties: time span active during a season (flowering duration for plants and foraging period for insect pollinators) and node degree (number of interaction partners), respectively. Plots of phenological sensitivity against time span and degree suggested weak positive relationships with both features for plants (Figs. 2a and 2c) and stronger positive relationships for pollinators (Figs. 2b and 2d).

**Fig. 2.**
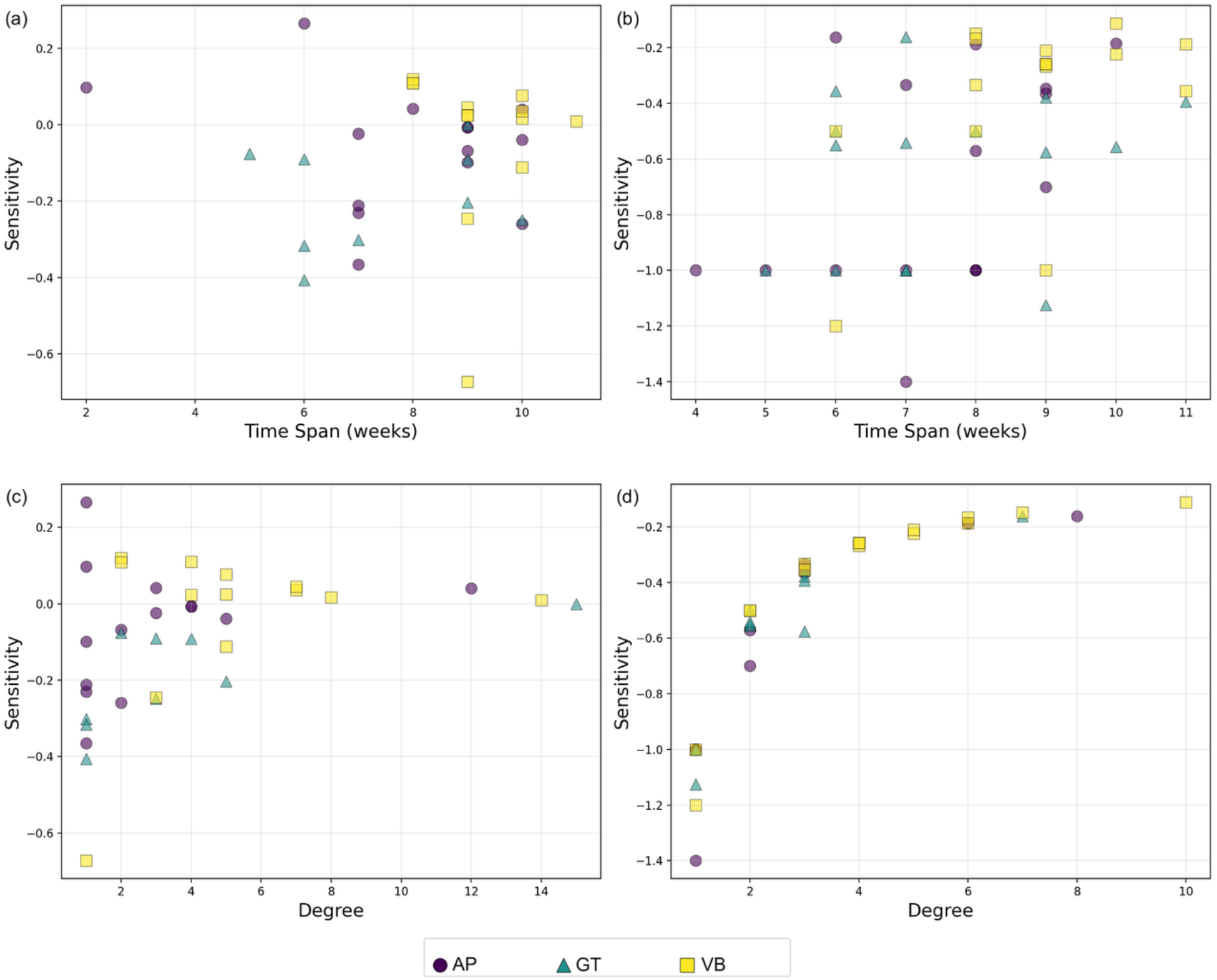
Factors influencing species-level sensitivity to phenology. Estimated relative changes in individual species densities for three field sites (“AP,” “GT,” and “VB”) following perturbations to temporal overlap with interaction partners; plant (left panels, a and c) and pollinator (right panels, b and d) sensitivity plotted against the length of time a species was active during a season (top panels, a and b) and its node degree, i.e., total number of observed interaction partners (bottom panels, c and d).

We conducted statistical analyses with mixed-effects models to evaluate the relationships in more detail. Models that included plant and pollinator sensitivity values together did not meet model assumptions, so we analyzed the two guilds separately (see Methods). We assessed models including time span, node degree, field site, and all combinations of interaction terms as fixed effects; when model assumptions were violated, we used the most-complex model that met model assumptions. For plants, this model included time span, node degree, field site and the interaction between time span and degree as fixed effects (Table 1). In this model, all three main effects were statistically significant, but the interaction term was not. For pollinators, following the non-linear relationship seen in Fig. 2d, we used inverse-transformed sensitivity values to ensure normality of residuals; the only model that met assumptions was the full model that included time span, node degree, field site, and all combinations of interaction terms as fixed effects (Table 2). In the pollinator-only model, time span as a main effect was not statistically significant, but degree was strongly significant. Site as a main effect was significant, as was its interaction with time span and degree separately, and there was a strongly significant three-way interaction between all three fixed effects.

**Table 1.** Analysis of Deviance Table (Type II Wald chi-square tests) for plants.

|  | Chi-sq | Df | p-value |  |
| --- | --- | --- | --- | --- |
| Time span | 4.54 | 1 | 0.033 | * |
| Degree | 11.85 | 1 | 0.00058 | *** |
| Site | 13.01 | 2 | 0.0015 | ** |
| Time span:Degree | 1.80 | 1 | 0.18 |  |

Significance codes: \*\*\* $p < 0.001$ , \*\* $p < 0.01$ , \* $p < 0.05$

**Table 2.**
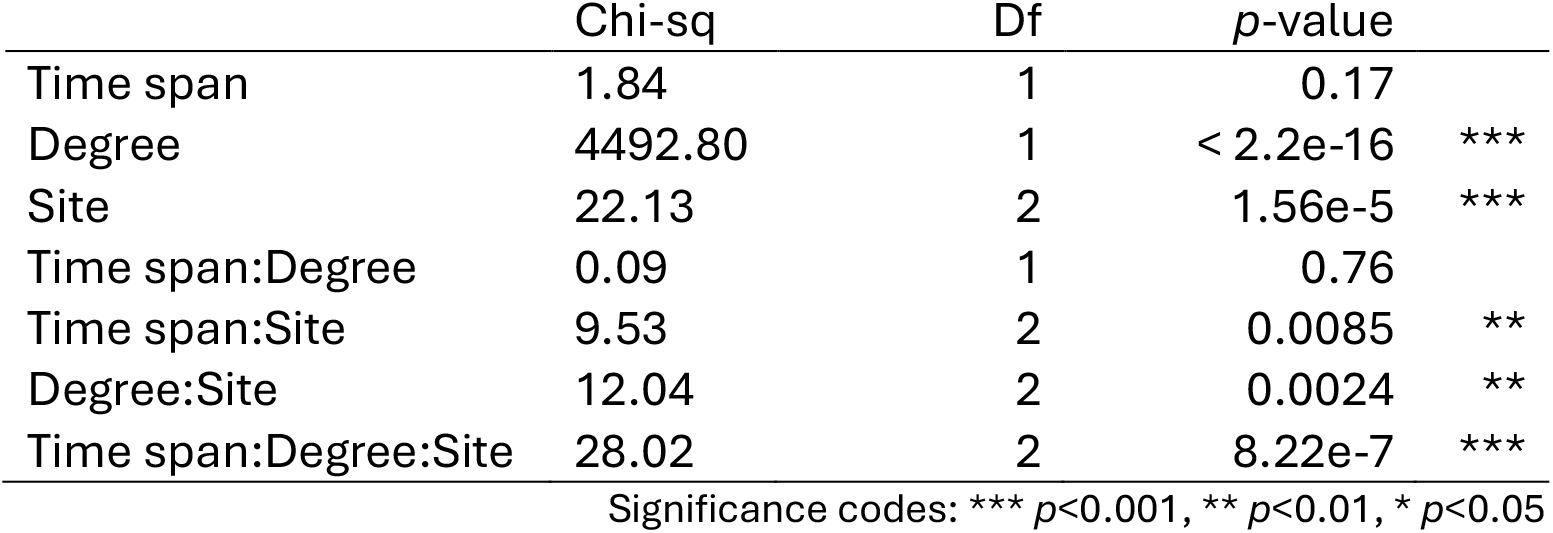
Analysis of Deviance Table (Type II Wald chi-square tests) for pollinators.

The regression models indicate that plants and pollinators with low degree (i.e., specialists) were most sensitive to decreases in temporal overlap with interaction partners, in line with our expectation that specialists are likely to be more sensitive to phenological shifts than generalists. Pollinators with shorter foraging periods were also more sensitive to decreases in temporal overlap, but with quantitatively different trends for the different sites.

## Implications for ecosystem resilience

Local stability analysis has been used for over fifty years as a standard framework for evaluating ecosystem responses to externally-driven perturbations. We showed that potential responses to changes to phenology are largely independent of traditional stability assessments made with respect to changes in species densities. Thus, ignoring phenology risks drawing misleading conclusions about a system’s inherent resilience to change. Our findings suggest that even very stable ecosystems, given just the right changes to interaction timing, could experience large deviations from long-term population densities.

The idea of “just the right changes” is worth further discussion. Our results refer to the initial impacts of phenological perturbations on population densities and do not account for subsequent effects. The precise trajectory of a system will be highly dependent on which species are affected by phenological changes and how important they (and their interaction partners) are to maintaining overall system stability. It is also worth noting that we considered *pulse perturbations* to established phenologies, that is, *temporary* changes. If phenological changes were to persist or intensify—as appears to be happening in nature (Inouye 2022)—then subsequent effects on species densities are likely to be amplified as the system transitions to a new dynamical state (Caravelli & Staniczenko 2016).

For the empirical plant-pollinator communities we analyzed, the strongest species-level result was that specialist pollinators were most sensitive to phenological perturbations. In some ways, this finding is to be expected: specialists receive benefits from a small set of interaction partners, so a change in the temporal availability of one of a specialist’s interaction partners is likely to have a disproportionately large effect on its population growth rate. The effect on generalists is less straightforward. Since generalists have many interaction partners, they are more likely to be affected by phenological changes to other species in the community, but their wider diet breadth means they may be better equipped to buffer phenological changes. The species-level sensitivity values used in our statistical analyses were averages over a species’ set of interaction partners, and closer inspection reveals substantial variability among sensitivity values within the interaction sets of some generalist (see Supplementary Materials). This means that the outcome of phenological changes affecting generalists, depending on which interactions are affected, can still be large. The topological structure of interactions is also important to consider. Our plant-pollinator networks displayed the common property of nestedness (Bascompte et al. 2003; Payrató-Borràs et al. 2020; Staniczenko & Panja 2023), in which specialists of one guild tend to interact with the generalists of the other guild. This means, for example, that multiple, vulnerable specialist pollinators are likely to be impacted by a change in phenology of a single generalist plant.

We focused on inter-guild mutualistic interactions, ignoring within-guild interactions such as competition among plants for pollination services and competition among pollinators for floral resources. Establishing the role of these competitive interactions in the context of interaction timing and ecosystem resilience will be revealing because phenology can function as an important life history strategy for reducing interspecific competition. As such, any shifts in phenological overlap will have direct implications for the competitive balance in a community and could lead to additional behavioral changes in response to environmental changes (e.g., interaction switching; Valdovinos et al. 2016; CaraDonna et al. 2017; Bartley et al. 2019).

## Temporal network data

Understanding how phenological changes will impact populations, species interactions, and ecosystem functions and services is a pressing question given accelerating environmental changes. A growing body of work demonstrates that phenological mismatches between interacting species are increasingly occurring in nature (Kudo et al. 2004; Thomson 2010; Diez et al. 2012; Kudo & Ida 2013; Theobald et al. 2017; Carter et al. 2018; Kehrberger & Holzschuh 2019; Duchenne et al. 2020), and both observational and experimental work suggest that such mismatches can have substantial negative impacts on the interacting partners (Parsche et al. 2011; Rafferty & Ives 2011; Gezon et al. 2016; Pardee et al. 2019; Gallagher & Campbell 2020; Waters et al. 2020). Nearly all these studies, however, focus on a single plant species and multiple pollinators, and work at the broader multi-species, community level (e.g., Rose-Person et al. 2024) is rare but needed.

To better understand the role of phenology in a community context, it is necessary to collect interaction data at a sufficiently fine temporal scale to capture phenological differences among species. We also recommend collecting multiple years of data to check for consistency in estimates of when species are active and their season-on-season population densities. Finally, it is valuable to compile datasets that include multiple, spatially independent sites to allow exploration of the generality of results.

Our approach can be applied to any ecological system for which phenologies are known or can be reliably estimated. We characterized this temporal information by a single new parameter, *ϕ*, which represents a mean-field approximation of the effects of interaction timing on population dynamics. This simplification makes it easier to collect suitable temporal data for parameterizing models and makes re-analyzing existing network data possible, but at the cost of averaging out subtle and potentially relevant intra-season fluctuations in interaction partner availability. Overall, our findings call for further work on the system-level impacts of phenological changes because they may deviate substantially from predictions based on existing community-based approaches and ecological theory.

## METHODS

### Field sites

Data were collected from 2016–2023 in three montane meadow sites (“AP,” “GT,” and “VB”), each separated by ∼2km, in the vicinity of the Rocky Mountain Biological Laboratory (“RMBL”), Colorado, USA (38°57.5′ N, 106°59.3′ W, ∼2900m above sea level).

### Data collection

Plant-pollinator network data and floral abundance data were sampled in each site weekly through the bulk of the growing season in these high-elevation sites, typically beginning in early June, shortly after snowmelt, and concluding in early August after most plant species had completed flowering. We observed plant-pollinator interactions in transects, with dedicated sampling time for each 1 × 2 m transect segment. On the same days that we collected network data in each site, we exhaustively censused flowers in our transects, recording species identities and floral counts.

#### Plant-pollinator visitation networks

We recorded plant-pollinator network data following Morozumi et al. 2022. Briefly, we sampled within four fixed 1m x 20m transects at each site. We divided transects into 2m-long sections and observed each section for 3 minutes per sample, divided into 90 seconds of observation time on each side of the transect, totaling 120 observation minutes per site per weekly sample. We paused observation timers while handling specimens and recording data to ensure the full sampling time was dedicated to observation. We randomized the sampling order of transect segments prior to each sampling event. This method contrasts with walking transects, where sampling time per area and per species is unknown, and in which visitor observability and floral abundance bias samples more strongly (e.g., Gibson et al. 2011). We only recorded interactions in which the flower visitor contacted the reproductive structures of the flower. We identified the flower visitor in the field to 30 gross taxonomic categories (used in this analysis) and collected the flower visitor wherever possible (except for bumble bees and butterflies, which we marked and released). We retained collected specimens for subsequent finer taxonomic identification. We randomly assigned observers to transect sections and randomized the order of transect segment sampled in each weekly sample to reduce biases. For subsequent data analyses, plant-pollinator visitation data were summed to create one network per site.

#### Plant and pollinator phenologies

We recorded plant and pollinator phenologies as the first and last date observed in each site in our data, for pollinators in the network data and for plants in our floral census (plants appearing in the network data were by design present in the census).

### Data analysis

[On publication, all code will be made freely available on GitHub.]

#### Parameterizing population dynamic models

For each site, empirical data specified binary network structure (location of non-zero elements in *α*), phenological coefficients (*ϕ*, calculated from first and last observations of each species across all years of data), and estimates of inter-season equilibrium species densities (using plant-visitation counts for pollinators and census data for plants). Given the lack of empirical values, intrinsic growth rates (*r*_*i*_ and *r*_*j*_), the cost to plants for providing resources to pollinators (*β*_*i*_), and half-saturation coefficients (*h*_*i*_, *h*_*j*_, and *e*_*i*_) were set to 1. Death rates (*d*_*i*_ and *d*_*j*_) were adjusted uniformly within each guild (i.e., all plant species had the same death rate, as did all pollinator species, but plants and pollinators had different death rates) with values chosen such that the system was always stable across all samples generated in the inverse problem approach, see below.

#### Inferring interaction strengths

These analyses were performed in Python (v3.11.8). We assumed that empirically recorded species densities represent inter-season equilibrium, or steady-state, conditions of the underlying population dynamic model (Eqn 1). Our implementation of the inverse problem approach meant finding interaction coefficients (*α*) that were consistent with these steady-state densities while also being constrained by fixed values for species-specific parameters (e.g., intrinsic growth and death rates). We also required interaction coefficients to be non-negative, reflecting mutualistic interactions between plants and pollinators. This formulation circumscribes an underdetermined system with many plausible combinations of interaction strengths (i.e., multiple plausible *α* matrices). We sampled from the feasible solution space using Artificial Centering Hit-and-Run (ACHR) (Kaufman & Smith 1998), a Markov chain Monte Carlo method. After a warm-up phase of 10,000 samples, we generated 1,000,000 samples and retained every 10th sample to reduce autocorrelation, yielding 100,000 approximately independent sets of interaction coefficients. For each sampled set of interaction coefficients (one *α* matrix), we computed the dominant eigenvalue from the corresponding community matrix to assess stability and computed the dominant singular value from the corresponding B matrix to assess phenological sensitivity.

#### Measuring ecosystem susceptibility to changes in phenology

Population dynamic models such as Eqn 1 can in general be represented as a system of coupled differential equations *dM*/*dt* = *f*(*M, ϕ*), where *M* is a vector of species densities, *ϕ* is a matrix of phenological overlaps, and *f*(*M, ϕ*) is a vector of linear or nonlinear functions. An equilibrium point is a non-negative vector *M\** and corresponding set of phenological overlaps *ϕ* = *ϕ\** such that *f*(*M*, ϕ\**) = 0; in this study, we estimated *M*=*M\** and *ϕ*=*ϕ\** from empirical data, as described above. The community matrix A is defined as A_lm_ = *∂*f_l_/*∂M*_*m*_|(*M*=*M*, ϕ*=*ϕ\**), which is the Jacobian matrix with partial derivatives calculated with respect to changes in species densities and evaluated at an equilibrium point (we use *l* and *m* as general indices for species to distinguish from *i* and *j* in the main text, which refer specifically to plant and pollinator species, respectively, in Eqn 1) (Levins 1968). An eigenvalue decomposition of the community matrix determines local stability, with the system considered stable if the real part of the dominant (largest) eigenvalue is negative. To assess ecosystem susceptibility to changes in phenology, partial derivatives are instead taken with respect to the phenology parameter, *ϕ*. The B matrix is defined as B_ln_ = -*∂f*_*l*_/*∂ϕ*_*n*_|(*M*=*M*, ϕ*=*ϕ\**), where *n* indexes interactions between species pairs. The minus sign indicates that we are considering the effect of a decrease in phenological overlap; a positive sign would represent an increase in phenological overlap, but this change only leads to a reversal in the predicted directional effect—increase or decrease in growth rate and therefore population density—following a perturbation, i.e., the magnitude of species’ response remains the same but the direction is switched. Since the B matrix is rectangular (species-by-interaction), the system-level susceptibility to changes in phenology is quantified using the dominant (largest) singular value, which is always positive. The larger the dominant singular value, the larger the collective first-order, initial response of species to a perturbation affecting phenology.

#### Species-level sensitivity regression models

These analyses were performed in the R statistical environment (v 4.4.0) using Linear Mixed Effects Models with the GLMMTMB package (Brooks et al. 2017; McGillycuddy et al. 2025). Model assumptions were assessed using the DHARMa package (Hartig 2024). We began with full models that included time span, node degree, field site, and all combinations of interaction terms as fixed effects. If model assumptions for the full model were violated, we sequentially deleted interaction terms, starting with site and then with degree, and retained the most-complex model that met model assumptions. Plant models met assumptions with no transformation of the response variable. Pollinator models did not, so we repeated the procedure starting with the full model but with different transformations of the response variable, including the Box-Cox family of transformations. Following this procedure, only one pollinator model met model assumptions: the full model with the response variable inverse-transformed. As multiple simpler plant models met assumptions, we validated our approach by comparing corrected AIC (AICc) values among the set of plant models that met assumptions; of those, the most-complex model (which we report on) also had the lowest AICc by >2. Finally, we compared the models we present with intercept-only null models using likelihood ratio tests. For both plants and pollinators, the non-null model had a significantly higher likelihood and substantially lower AICc (≫2) in both cases.

## Author Contributions

P.P.A.S. and D.P. designed research; B.J.B. collected data; J.V. performed modeling analyses; B.J.B. performed statistical analyses; P.P.A.S. and J.V. wrote the first draft; and P.P.A.S., J.V., B.J.B. and D.P. contributed to subsequent drafts.

## Acknowledgements

The authors thank Julia Amoroso, Rampal S. Etienne, Anje-Margriet Neutel, and Mark Novak for comments on earlier drafts of the manuscript.

## REFERENCES

Allesina, S. & Tang, S. (2012). Stability criteria for complex ecosystems. Nature 483, 205–208. 10.1038/nature10832

Allesina, S. & Tang, S. (2015). The stability-complexity relationship at age 40: A random matrix perspective. Popul. Ecol. 57, 63–75. 10.1007/s10144-014-0471-0

Anderson, C. R., Curtsdotter, A. R. K., Staniczenko, P. P. A., Valdovinos, F. S. & Brosi, B. J. (2024). The interplay of binary and quantitative structure on the stability of mutualistic networks. Integrative and Comparative Biology 64, 827–840. 10.1093/icb/icae074

Ashman, T.-L. et al. (2004). Pollen limitation of plant reproduction: ecological and evolutionary causes and consequences. Ecology 85, 2408–2421. 10.1890/03-8024

Bartley, T. J. et al. (2019). Food web rewiring in a changing world. Nat. Ecol. Evol. 3, 345–354. 10.1038/s41559-018-0772-3

Bartomeus, I. et al. (2011). Climate-associated phenological advances in bee pollinators and bee-pollinated plants. Proc. Natl Acad. Sci. USA 108, 20645–20649. 10.1073/pnas.1115559108

Bascompte, J., Jordano, P., Melián, C. J. & Olesen, J. M. (2003). The nested assembly of plant–animal mutualistic networks. Proc. Natl Acad. Sci. USA 100, 9383–9387. 10.1073/pnas.1633576100

Brooks, et al. (2017). glmmTMB Balances Speed and Flexibility Among Packages for Zero-inflated Generalized Linear Mixed Modeling. The R Journal 9, 378–400. 10.32614/RJ-2017-066

Brose, U., Williams, R. J. & Martinez, N. D. (2006). Allometric scaling enhances stability in complex food webs. Ecol. Lett. 9, 1228–1236. 10.1111/j.1461-0248.2006.00978.x

Brosi, B. J. (2016). Pollinator specialization: from the individual to the community. New Phytol. 210, 1190–1194. 10.1111/nph.13951

CaraDonna, P. J., Petry, W. K., Brennan, R. M., Cunningham, J. L., Bronstein, J. L., Waser, N. M. & Sanders, N. J. (2017). Interaction rewiring and the rapid turnover of plant-pollinator networks. Ecol. Lett. 20, 385–394. 10.1111/ele.12740

Caravelli, F. & Staniczenko, P. P. A. (2016). Bounds on transient instability for complex ecosystems. PLOS ONE 11, e0157876. 10.1371/journal.pone.0157876

Carter, S. K., Saenz, D. & Rudolf, V. H. W. (2018). Shifts in phenological distributions reshape interaction potential in natural communities. Ecol. Lett. 21, 1143–115. 10.1111/ele.13081

Diez, J. M. et al. (2012). Forecasting phenology: from species variability to community patterns. Ecol. Lett. 15, 545–553. 10.1111/j.1461-0248.2012.01765.x

Duchenne, F., Thébault, E., Michez, D. et al. (2020). Phenological shifts alter the seasonal structure of pollinator assemblages in Europe. Nat. Ecol. Evol. 4, 115–121. 10.1038/s41559-019-1062-4

Duchenne, F., Wüest, R. O. & Graham, C. H. (2022). Seasonal structure of interactions enhances multidimensional stability of mutualistic networks. Proc. R. Soc. B 289, 20220064. 10.1098/rspb.2022.0064

Gallagher, M. K. & Campbell, D. R. (2020). Pollinator visitation rate and effectiveness vary with flowering phenology. Am. J. Bot. 107, 445–455. 10.1002/ajb2.1439

Gellner, G. & McCann, K. S. (2016). Consistent role of weak and strong interactions in high-and low-diversity trophic food webs. Nat. Commun. 7, 11180. 10.1038/ncomms11180

Gellner, G., McCann, K. S. & Hastings, A. (2023). Stable diverse food webs become more common when interactions are more biologically constrained, Proc. Natl. Acad. Sci. USA 120, e2212061120. 10.1073/pnas.2212061120

Gezon, Z. J., Inouye, D. W. & Irwin, R. E. (2016). Phenological change in a spring ephemeral: implications for pollination and plant reproduction. Glob. Change Biol. 22, 1779–1793. 10.1111/gcb.13209

Gibson, R. H., Knott, B., Eberlein, T. & Memmott, J. (2011). Sampling method influences the structure of plant-pollinator networks. Oikos 120, 822–831. 10.1111/j.1600-0706.2010.18927.x

Glaum, P., Wood, T.J., Morris, J.R. & Valdovinos, F.S. (2021). Phenology and flowering overlap drive specialisation in plant-pollinator networks. Ecol. Lett. 24, 2648–2659. 10.1111/ele.13884

Golub, G. H. & Van Loan, C. F. (1996). Matrix Computations (3rd ed.), Section 8.8.6. Johns Hopkins.

Hartig, F. (2024). DHARMa: Residual Diagnostics for Hierarchical (Multi-Level / Mixed) Regression Models. R package version 0.4.7. https://CRAN.R-project.org/packageffDHARMa

Holland, J. N. & DeAngelis, D. L. (2010). A consumer-resource approach to the density-dependent population dynamics of mutualism. Ecology 91, 1286–1295. 10.1890/09-1163.1

Inouye, D. W. (2022). Climate change and phenology. WIREs Climate Change 13, e764. 10.1002/wcc.764

Jacquet, C., Moritz, C., Morissette, L. et al. (2016). No complexity-stability relationship in empirical ecosystems. Nat. Commun. 7, 12573. 10.1038/ncomms12573

Kaufman, D. E. & Smith, R. L. (1998). Direction Choice for Accelerated Convergence in Hit-and-Run Sampling. Oper. Res. 46, 84–95. http://www.jstor.org/stable/223065

Kehrberger, S. & Holzschuh, A. (2019). Warmer temperatures advance flowering in a spring plant more strongly than emergence of two solitary spring bee species. PLoS ONE 14, e0218824. 10.1371/journal.pone.0218824

Kudo, G., Nishikawa, Y., Kasagi, T. & Kosuge, S. (2004). Does seed production of spring ephemerals decrease when spring comes early? Ecol. Res. 19, 255–259. 10.1111/j.1440-1703.2003.00630.x

Kudo, G. & Ida, T. Y. (2013). Early onset of spring increases the phenological mismatch between plants and pollinators. Ecology 94, 2311–2320. 10.1890/12-2003.1

Levins, R. (1968). Evolution in Changing Environments: Some Theoretical Explorations. Princeton University Press, Princeton.

May, R. M. (1972). Will a large complex system be stable? Nature 238, 413–414. 10.1038/238413a0

May, R. M. (1973). Stability and complexity in model ecosystems. Princeton University Press, Princeton.

McCann, K. S. (2000). The diversity-stability debate. Nature 405, 228–233. 10.1038/35012234

McGillycuddy, M., Popovic, G., Bolker, B. M., & Warton, D. I. (2025). Parsimoniously Fitting Large Multivariate Random Effects in glmmTMB. Journal of Statistical Software 112, 1–19. 10.18637/jss.v112.i01

Memmott, J., Craze, P. G., Waser, N. M. & Price, M. V. (2007). Global warming and the disruption of plant-pollinator interactions. Ecol. Lett. 10, 710–717. 10.1111/j.1461-0248.2007.01061.x

Miller-Rushing, A. J., Høye, T. T., Inouye, D. W. & Post, E. (2010). The effects of phenological mismatches on demography. Philos. Trans. R. Soc. Lond. B. Biol. Sci. 365, 3177–3186. 10.1098/rstb.2010.0148

Morozumi, C., Loy, X., Reynolds, V., Schiffer, A., Morrison, B., Savage, J. and Brosi, B. J. (2022). Simultaneous niche expansion and contraction in plant-pollinator networks under drought. Oikos e09265. 10.1111/oik.09265

Neutel, A.-M. et al. (2007). Reconciling complexity with stability in naturally assembling food webs. Nature 449, 599–603. 10.1038/nature06154

Ovaskainen, O. et al. (2013). Community-level phenological response to climate change. Proc. Natl. Acad. Sci. U.S.A. 110, 13434–13439, 10.1073/pnas.1305533110

Pardee, G.L., Jensen, I.O., Inouye, D.W. & Irwin, R.E. (2019). The individual and combined effects of snowmelt timing and frost exposure on the reproductive success of montane forbs. J. Ecol. 107, 1970–1981. 10.1111/1365-2745.13152

Parsche, S., Fründ, J. & Tscharntke, T. (2011). Experimental environmental change and mutualistic vs. antagonistic plant flower-visitor interactions. Perspect. Pl. Ecol. Evol. Syst. 13, 27–35. 10.1016/j.ppees.2010.12.001

Parmesan, C. (2007). Influences of species, latitudes and methodologies on estimates of phenological response to global warming. Glob. Change Biol. 13, 1860–1872. 10.1111/j.1365-2486.2007.01404.x

Payrató-Borràs, C., Hernández, L. & Moreno Y. (2020). Measuring nestedness: A comparative study of the performance of different metrics. Ecol. Evol. 10: 11906–11921. 10.1002/ece3.6663

Peng, S., Ellison, A. M. & Davis, C. C. (2025). Climate change intensifies plant-pollinator mismatch and increases secondary extinction risk for plants in northern latitudes. Proc. Natl. Acad. Sci. U.S.A. 122, e2506265122. 10.1073/pnas.2506265122

Peralta, G., Vázquez, D. P., Chacoff, N. P., Lomáscolo, S. B., Perry, G. L. W. & Tylianakis, J. M. (2020). Trait matching and phenological overlap increase the spatio-temporal stability and functionality of plant–pollinator interactions. Ecol. Lett. 23, 1107–1116. 10.1111/ele.13510.704

Peralta, G. et al. (2024). Predicting plant–pollinator interactions: concepts, methods, and challenges. Trends Ecol. Evol., 39, 494–505. 10.1016/j.tree.2023.12.005.700

R Core Team (2021). R: A language and environment for statistical computing. R Foundation for Statistical Computing, Vienna, Austria. https://www.R-project.org/

Rafferty, N. E. & Ives, A. R. (2011). Effects of experimental shifts in flowering phenology on plant-pollinator interactions. Ecol. Lett. 14, 69–74. 10.1111/j.1461-0248.2010.01557.x

Ramos-Jiliberto, R., Moisset de Espanés, P., Franco-Cisterna, M., Petanidou, T. & Vázquez, D. P. (2018). Phenology determines the robustness of plant-pollinator networks. Sci. Rep. 8, 14873. 10.1038/s41598-018-33265-6

de Ruiter, P. C., Neutel, A.-M. & Moore, J. C. (1995). Energetics, patterns of interaction strengths and stability in real ecosystems. Science 269, 1257–1260. 10.1126/science.269.5228.1257

Rose-Person, A., Spasojevic, M. J., Forrester, C. et al. (2024). Early snowmelt advances flowering phenology and disrupts the drivers of pollinator visitation in an alpine ecosystem. Alp. Botany 134, 141–150. 10.1007/s00035-024-00315-x

Roslin, T., Antão, L., Hällfors, M. et al. (2021) Phenological shifts of abiotic events, producers and consumers across a continent. Nat. Clim. Chang. 11, 241–248. 10.1038/s41558-020-00967-7

Staniczenko, P. P. A. & Panja, D. (2023). Temporal origin of nestedness in interaction networks. PNAS Nexus 2, pgad412. 10.1093/pnasnexus/pgad412

Stone, L. (2020). The stability of mutualism. Nat. Commun. 11, 2648. 10.1038/s41467-020-16474-4

Tang, S. & Allesina, S. (2014). Reactivity and stability of large ecosystems. Front. Ecol. Evol. 2, 21. 10.3389/fevo.2014.00021

Thackeray. S. J, et al. (2016). Phenological sensitivity to climate across taxa and trophic levels. Nature 535, 241–255. 10.1038/nature18608

Theobald, E. J., Breckheimer, I. & HilleRisLambers, J. (2017). Climate drives phenological reassembly of a mountain wildflower meadow community. Ecology 98, 2799–2812. 10.1002/ecy.1996

Thomson, J. D. (2010). Flowering phenology, fruiting success and progressive deterioration of pollination in an early-flowering geophyte. Philos. Trans. R. Soc. Lond. B Biol. Sci. 365, 3187–3199. 10.1098/rstb.2010.0115

Valdovinos, F. S., Brosi, B. J., Briggs, H. M., de Espanés, P. M., Ramos-Jiliberto, R. & Martinez, N. D. (2016). Niche partitioning due to adaptive foraging reverses effects of nestedness and connectance on pollination network stability. Ecol. Lett. 19, 1277–1286. 10.1111/ele.12664

Waters, S.M., Chen, W.-L. C., HilleRisLambers, J. (2020). Experimental shifts in exotic flowering phenology produce strong indirect effects on native plant reproductive success. J. Ecol. 108, 2444–2455. 10.1111/1365-2745.13392

Wilson, D. S. & Yoshimura, J. (1994). On the coexistence of specialists and generalists. Am. Nat. 144, 692–707. http://www.jstor.org/stable/2462946

