## Supplementary Materials for "Susceptibility of ecosystems to interaction timing"

### Supplementary Materials
### Susceptibility of ecosystems to interaction timing

Phillip P. A. Staniczenko[1,2,3,#,*], Jilles Verwoerd[4,#], Berry J. Brosi[5,6], Debabrata Panja[4,*]

[1]Department of Biology, Brooklyn College, City University of New York, Brooklyn NY 11210, USA
[2]Biology PhD Program, CUNY Graduate Center, New York NY 10016, USA
[3]Division of Invertebrate Zoology, American Museum of Natural History, New York NY 10024, USA
[4]Department of Information and Computing Sciences, Utrecht University, Utrecht 3584 CC, The Netherlands
[5]Department of Biology, University of Washington, Seattle WA 98195, USA
[6]Rocky Mountain Biological Laboratory, Crested Butte CO 81224, USA

#### Contents

**A: Data** **2**

**B: Data processing** **2**

**C: Model** **2**

**D: Parameters** **4**

### A: Data

This study uses two types of dataset: (1) interaction data on plant visits by insect pollinators and (2) plant abundance data. For additional details on the data sampling protocol, see [1].

Data were collected in three montane meadow sites (Avery Picnic, Gothic Town, Virginia Basin), each separated by  2km, in the vicinity of the Rocky Mountain Biological Laboratory ("RMBL"), Colorado, USA (38°57.5′ N, 106°59.3′ W, ∼2900m above sea level). Collection took place during the flowering season (July–August). Sites span a range of elevations but share similar latitudes and longitudes. At each site, transects were surveyed weekly during the flowering season. During each visit, field ecologists observed the transects for a fixed time interval, recording all interactions between pollinators and the reproductive parts of flowering plants that occurred during that period. Insects involved in interactions were captured and labeled for species identification. Each recorded interaction includes the date, time, location, and identities of both plant and pollinator species.

Flower abundance data were collected by counting open flowers along the same transects throughout the season. No comparable census data were collected for pollinators due to their mobility.

### B: Data processing

We derived three key quantities from the data for each site: (1) interaction network structure, (2) relative species densities, and (3) phenological overlap coefficients.

1. For each site, we constructed a bipartite network representing plant-pollinator interactions aggregated across the eight years (2016–2023). Nodes represent species and edges connect pollinators to the plants they visited. We included only species observed in at least 6 of the 8 years, and only retained interactions that were observed at least 3 times. This process resulted in a 14 plant species $\times$ 16 pollinator species network with 41 interspecific interactions for the Avery Picnic site, a 9 $\times$ 16 network with 35 interactions for the Gothic Town site, and a $13 \times 16$ network with 67 interactions for the Virginia Basin site.

2. We estimated relative species densities for both plants and pollinators. For pollinators, we used interaction frequency as a proxy for abundance, assuming that more frequently observed species are more common. We summed the total number of interactions for each pollinator across all years and divided each by the highest total among pollinators, thereby normalizing densities between zero and unity. For plants, we used the census data, averaging flower counts across the eight years and normalizing each average by dividing it by the highest average observed among all plant species.

3. To capture the effect of seasonal activity on interactions, we computed phenological overlap coefficients $\phi_{ij} \in [0, 1]$ that scale interaction strength based on overlap in activity periods (see below and Eqn 1 in the main text). For each species $i$, we defined an activity window $I_i = [\tau_i^{(0)}, \tau_i^{(1)}]$, where $\tau_i^{(0)}$ is the time it becomes active and $\tau_i^{(1)}$ is the time it becomes dormant. These dates were extracted from the first and last observations across all years, using interaction data for pollinators and the extreme values out of interaction data and census data for plants. The overlap between two species' activity windows is given by $|I_i \cap I_j|$, and the phenological overlap coefficient is defined as

$$\phi_{ij} = \frac{|I_i \cap I_j|}{|I_i|} \tag{1}$$

This measures the proportion of species $i$'s active period that overlaps with species $j$'s availability. Since the overlap is normalized relative to $i$, the coefficients are generally asymmetric: $\phi_{ij} \neq \phi_{ji}$.

### C: Model

This study considers the general nonlinear model

$$\frac{dM_i}{dt} = f_i(M_i) + \sum_{j \in S} \alpha_{ij} g_{ij}(M_i M_j) \tag{2}$$

where $S$ is the set of species in the network, and $\alpha_{ij}$ represents the interaction coefficients between species $i$ and $j$. If two species do not interact, their corresponding $\alpha_{ij}$ is set to zero.

The key features of this model are: (1) linearity in the interaction coefficients $\alpha_{ij}$, and (2) interaction effects $g_{ij}(M_i M_j)$ that are proportional to the product of species abundances $M_i M_j$.

To incorporate temporal constraints, we introduce coefficients that scale the product of abundances to reflect the phenological overlap of species throughout the season. This modification allows the model to account for how timing influences species interactions, leading to the general model

$$\frac{dM_i}{dt} = f_i(M_i) + \sum_{j \in S} \phi_{ij} \alpha_{ij} g_{ij}(M_i M_j) \tag{3}$$

While our approach applies broadly, here we focus specifically on plant-pollinator networks. We build on the well-established model by Holland & DeAngelis [2]. This model captures unidirectional consumer-resource mutualism, where in addition to species of both guilds benefiting from an interaction, pollinators additionally act as consumers and plants as resources, reflecting how plants provide nectar in exchange for pollination services.

Building on the Holland-DeAngelis framework, we model the population dynamics of plants and pollinators by the following system of differential equations:

$$\frac{dM_i}{dt} = r_i M_i + \sum_{j \in S_A} \alpha_{ij} \frac{\phi_{ij} M_i N_j}{h_j + N_j} - \beta_i \sum_{j \in S_A} \frac{\phi_{ij} M_i N_j}{e_i + M_i} - d_i M_i^2 \tag{4}$$

$$\frac{dN_j}{dt} = r_j N_j + \sum_{i \in S_P} \alpha_{ij} \frac{\phi_{ji} M_i N_j}{h_i + M_i} - d_j N_j^2 \tag{5}$$

The first equation describes the population dynamics of plant densities $M_i$, while the second describes pollinator densities $N_j$. A table of the variables and parameters used in the model is given below:

- $S_A$: Set of all pollinator species in the network
- $S_P$: Set of all plant species in the network
- $M_i$: Density of plant species $i$
- $N_j$: Density of pollinator species $j$
- $\alpha_{ij}$: Interaction coefficient between species $i$ and $j$, where $\alpha_{ij} = 0$ if two species do not interact
- $\phi_{ij}$: Phenological overlap coefficient
- $r_i$, $r_j$: Intrinsic growth rate of species $i$, $j$
- $d_i$, $d_j$: Intrinsic death rate of species $i$, $j$
- $h_i$, $h_j$: Half-saturation constant for species $i$, $j$
- $\beta_i$: Cost to plant species $i$ for producing resources to attract pollinators
- $e_i$: Half-saturation constant for species $i$

We extended the Holland-DeAngelis model by explicitly incorporating the phenological overlap coefficients $\phi_{ij} \in [0, 1]$. Ignoring phenology corresponds to setting $\phi_{ij} = 1$ for all interactions. These coefficients scale down the contribution of interactions to species population dynamics. The effects of this extension are twofold: (1) the overall contribution of interactions to species density dynamics decreases and (2) the distribution of the $\phi_{ij}$ for species $i$ is weighted towards the species $j$ with which it shares the greatest overlap.

For the terms involving saturation constants in Eqns. 4 and 5, the functional forms assume that the benefits and costs of each interspecific interaction are not equivalent and are therefore not substitutable with one another. An alternative functional form is, e.g., $\frac{M_i \sum_{j \in S_A} \alpha_{ij} \phi_{ij} N_j}{h_i + \sum_{j \in S_A} \alpha_{ij} \phi_{ij} N_j}$, which assumes that interspecific interactions are substitutable, with benefits or costs saturating from the perspective of the focal species, rather scaling distinctly for each interaction partner as in Eqns. 4 and 5. We do not expect the choice of saturation functional form to change results qualitatively given that local stability analysis assumes empirical systems are observed at equilibrium states, with unchanging season-on-season maximum population densities.

The model describes species density dynamics on a seasonal (yearly) time-scale, tracking changes in plant and pollinator populations across successive flowering seasons.

### D: Parameters

While parameters such as intrinsic growth rates $r_i$ could be measured experimentally, this process is often difficult and time-consuming. Therefore, we choose reasonable values for these parameters to focus on the model's behavior and stability.

We set intrinsic parameters $r_i$, $r_j$, $e_i$, $h_i$, $h_j$, and $\beta_i$ to 1. The death rates $d_i$ and $d_j$ are chosen to ensure two critical conditions are met: (1) **feasibility**, a steady-state exists with biologically realistic non-negative interaction coefficients $\alpha_{ij} \geq 0$, and (2) **self-regulation**, diagonal elements of the Jacobian are negative, preventing unbounded population growth.

##### Feasibility Condition

The feasibility condition ensures that the sampling space for interaction coefficients $\alpha$ is non-empty. For pollinators, from the steady-state equation we require

$$r_j - d_j N_j < 0 \implies d_j > \frac{r_j}{N_j} \tag{6}$$

For plants, the feasibility condition additionally accounts for the cost term in the dynamics, i.e.,

$$r_i - d_i M_i - \frac{\beta_i}{(e_i + M_i)} \sum_j \phi_{ij} N_j < 0 \implies d_i > \frac{r_i - \frac{\beta_i}{(e_i+M_i)} \sum_j \phi_{ij} N_j}{M_i} \tag{7}$$

These bounds differ because pollinators and plants have different governing differential equations, Eqns. 4 and 5, respectively. The cost term $\beta_i$ in the plant equation introduces additional complexity in the feasibility constraint.

##### Self-regulation Condition

The self-regulation condition ensures negative diagonal elements in the Jacobian matrix at steady-state, which corresponds to negative feedback from intraspecific competition. For pollinators, this requires

$$-d_j N_j < 0 \implies d_j > 0 \tag{8}$$

For plants, the self-regulation condition must account for the derivative of the cost term and so

$$\frac{\beta_i}{(e_i + M_i)^2} \sum_j \phi_{ij} M_i N_j - d_i M_i < 0 \implies d_i > \frac{\beta_i}{(e_i + M_i)^2} \frac{1}{M_i} \sum_j \phi_{ij} M_i N_j \tag{9}$$

The difference again stems from the cost term in the plant dynamics, which contributes to the diagonal elements of the Jacobian.

##### Combined conditions

To satisfy both feasibility and self-regulation simultaneously, we set $d_i$ to the maximum of the two bounds. For pollinators

$$d_j = \max \left\{ 0, \frac{r_j}{N_j} \right\} = \frac{r_j}{N_j} \tag{10}$$

and for plants

$$d_i = \max \left\{ \frac{\beta_i}{(e_i + M_i)^2} \frac{1}{M_i} \sum_j \phi_{ij} M_i N_j, \frac{r_i - \frac{\beta_i}{(e_i+M_i)} \sum_j \phi_{ij} N_j}{M_i} \right\} \tag{11}$$

##### Tuning parameters

To control the strength of these conditions and explore different stability regimes, we introduce tuning hyperparameters $\varepsilon_f < 0$ and $\varepsilon_s < 0$.

For the feasibility condition, we introduce $\varepsilon_f$ such that for pollinators

$$r_j - d_j N_j = \varepsilon_f \implies d_j = \frac{r_j - \varepsilon_f}{N_j} \tag{12}$$

and for plants

$$r_i - d_i M_i - \frac{\beta_i}{(e_i + M_i)} \sum_j \phi_{ij} N_j - \varepsilon_f = 0 \implies d_i = \frac{r_i - \frac{\beta_i}{(e_i+M_i)} \sum_j \phi_{ij} N_j - \varepsilon_f}{M_i} \tag{13}$$

For the self-regulation condition, we introduce $\varepsilon_s$ for plants:

$$\frac{\beta_i}{(e_i + M_i)^2} \sum_j \phi_{ij} M_i N_j - d_i M_i - \varepsilon_s = 0 \implies d_i = \frac{1}{M_i} \left( \frac{\beta_i}{(e_i + M_i)^2} \sum_j \phi_{ij} M_i N_j - \varepsilon_s \right) \tag{14}$$

With these tuning parameters, the final death rates are:

$$d_j = \frac{r_j - \varepsilon_f}{N_j} \quad \text{for pollinators and} \tag{15}$$

$$d_i = \max \left\{ \frac{1}{M_i} \left( \frac{\beta_i}{(e_i + M_i)^2} \sum_j \phi_{ij} M_i N_j - \varepsilon_s \right), \frac{r_i - \frac{\beta_i}{(e_i+M_i)} \sum_j \phi_{ij} N_j - \varepsilon_f}{M_i} \right\} \quad \text{for plants.} \tag{16}$$

As $\varepsilon_f \to 0$, feasibility constraints tighten, making it harder to find valid steady-states with non-negative interaction coefficients. As $\varepsilon_s \to 0$, self-regulation weakens, leading to reduced stability and potential for unbounded growth. By tuning $\varepsilon_f$ and $\varepsilon_s$, we can explore different stability regimes while maintaining biological realism. Importantly, once these hyperparameters are set, the death rates $d_i$ and $d_j$ are automatically determined to satisfy both conditions.

### E: The inverse problem

In ecological modeling, interaction coefficients $\alpha_{ij}$ are typically treated as known inputs used to predict species densities as outputs. However, directly measuring interaction coefficients in the field is challenging. By contrast, relative species densities can be estimated more readily through field observations.

The inverse problem offers a practical alternative: we use empirically observed (estimates of) species densities as inputs to infer the corresponding interaction coefficients, $\alpha_{ij}$. This approach relies on a key assumption: observed seasonal maxima of species densities reflect stable steady-states of the underlying system.

#### Mathematical formulation

The steady-state equations at observed densities $M_i^*$ and $N_j^*$ are

$$\left.\frac{dM_i}{dt}\right|_{M_i^*, N_j^*} = r_i M_i^* + \sum_{j \in S_A} \phi_{ij} \alpha_{ij} \frac{M_i^* N_j^*}{h_j + N_j^*} - \beta_i \sum_{j \in S_A} \phi_{ij} \frac{M_i^* N_j^*}{e_i + M_i^*} - d_i (M_i^*)^2 = 0 \tag{17}$$

$$\left.\frac{dN_j}{dt}\right|_{M_i^*, N_j^*} = r_j N_j^* + \sum_{i \in S_P} \phi_{ji} \alpha_{ij} \frac{M_i^* N_j^*}{h_i + M_i^*} - d_j (N_j^*)^2 = 0 \tag{18}$$

A crucial observation is that while these equations are nonlinear in species densities, they are *linear* in the interaction coefficients $\alpha_{ij}$. This linearity makes the inverse problem computationally tractable for large ecological networks. Furthermore, each $\alpha_{ij}$ appears in only one equation corresponding to species $i$, allowing the system to be decomposed into independent subproblems for each species.

#### Linear programming formulation

By requiring $\alpha_{ij} \geq 0$ to reflect mutualistic plant-pollinator interactions, the inverse problem becomes a linear programming problem. For each plant species $i$, we solve for the following conditions:

$$\text{find } \alpha_{ij} \geq 0 \text{ such that } r_i M_i^* + \sum_{j \in S_A} \phi_{ij} \alpha_{ij} \frac{M_i^* N_j^*}{h_j + N_j^*} - \beta_i \sum_{j \in S_A} \phi_{ij} \frac{M_i^* N_j^*}{e_i + M_i^*} - d_i (M_i^*)^2 = 0 \tag{19}$$

and similarly, for each pollinator species $j$:

$$\text{find } \alpha_{ij} \geq 0 \text{ such that } r_j N_j^* + \sum_{i \in S_P} \phi_{ji} \alpha_{ij} \frac{M_i^* N_j^*}{h_i + M_i^*} - d_j (N_j^*)^2 = 0 \tag{20}$$

Standard linear programming solvers can efficiently find solutions satisfying these constraints, or determine that no feasible solution exists.

##### Stability analysis

To verify whether the inferred coefficients correspond to a stable steady-state, we compute the Jacobian matrix $J_\alpha$ evaluated at $(M_i^*, N_j^*)$, commonly known as the community matrix [3], and examine its dominant eigenvalue (the eigenvalue with the largest real part). A negative real part of the dominant eigenvalue, $\lambda_{\max} < 0$, indicates local asymptotic stability.

If the system is found to be unstable ($\lambda_{\max} \geq 0$), we adjust the tuning parameters $\varepsilon_f$ and $\varepsilon_s$ to modify the death rates $d_i$ until stability is just achieved (i.e., marginal stability). This iterative approach ensures that the inferred interaction coefficients are consistent with both the observed species densities and ecological stability requirements.

#### F: Sampling interaction coefficients

##### An underdetermined system

In complex ecosystems, interactions outnumber species. Each interaction is characterized by a coefficient $\alpha_{ij}$, while each species contributes only one steady-state equation. This leads to an underdetermined system with more unknowns than equations. Consequently, if a feasible solution exists, it is typically not unique—there exists a high-dimensional polytope of valid interaction coefficients consistent with the observed steady-state densities.

To address this, we frame the inverse problem statistically by sampling from the space of feasible solutions. This allows us to assess the robustness of ecological properties (such as stability and phenological sensitivity) across the range of possible interaction sets of interaction coefficients.

##### Hit-and-run sampling

The feasible region forms a convex polytope defined by the linear constraints from the steady-state equations and the non-negativity requirements $\alpha_{ij} \geq 0$. To sample uniformly from this polytope, we employ Artificial Centering Hit-and-Run (ACHR) [4], a Markov chain Monte Carlo method well-suited for high-dimensional convex spaces.

Standard Hit-and-Run sampling can be inefficient in spaces with irregular geometry, such as elongated feasible regions that are narrow along some dimensions and wide along others. This structure commonly arises in ecological systems in which certain interactions are tightly constrained while others remain flexible. ACHR addresses this limitation by biasing sampling directions toward less constrained regions. Specifically, on each iteration, the algorithm:

1. Selects a random direction from the set of vectors pointing from previous sample points to an artificial center (the running mean of previous samples).
2. Moves along this direction to a new point within the feasible region.
3. Updates the artificial center with the new sample.

Since directions allowing longer steps naturally shift the mean more significantly, the algorithm favors these directions, enhancing sampling efficiency and coverage of the solution space.

##### Finding an interior point

ACHR requires an initial strictly interior point within the feasible region. Finding such a point in high-dimensional spaces is nontrivial. We employ a log-barrier method combined with a penalty function to locate a point near the center of the polytope.

The objective function we maximize is

$$f(\boldsymbol{\alpha}) = \sum_k [\log(\alpha_k - \ell_k) + \log(u_k - \alpha_k)] - \sum_k \log\left(e^{\gamma(\alpha_k - \ell_k)} + e^{\gamma(u_k - \alpha_k)}\right) \tag{21}$$

where $\ell_k = 0 + \epsilon$ and $u_k = 1000 - \epsilon$ are the strictly interior bounds on $\alpha_k$ (with $\epsilon = 10^{-6}$), and $\gamma = 10$ is a penalty parameter. The log-barrier terms push the solution away from the boundaries, while the penalty term (a smooth approximation of the minimum distance to boundaries) encourages centrality. We solve this optimization problem using the Clarabel solver [5] with feasibility tolerance set to $10^{-10}$.

##### Sampling protocol

Our sampling procedure consists of the following steps:

1. **Warm-up phase**: Generate and discard 10,000 initial samples to allow the Markov chain to reach its stationary distribution and eliminate bias from the starting point.
2. **Production phase**: Generate 1,000,000 samples from the feasible region using ACHR.
3. **Thinning**: Retain every 10th sample to reduce serial correlation between consecutive samples, following [4]. This yields 100,000 approximately independent samples.
4. **Analysis**: For each retained sample of interaction coefficients $\{\alpha_{ij}\}$, we compute:
   - The dominant eigenvalue $\lambda_{\max}$ of the community matrix to assess stability.
   - The phenological sensitivity matrix, referred to as the "$B$ matrix" in the main text (described in the following section) to quantify species responses to perturbations in phenology.

This protocol ensures unbiased and representative coverage of the solution space while maintaining computational efficiency. The large sample size allows us to reliably characterize the distribution of stability and sensitivity properties across all feasible interaction configurations.

#### G: Stability analysis

For each set of sampled interaction coefficients, we assess local stability by taking partial derivatives of our dynamical system (Eqns. 4 and 5) with respect to species densities and then evaluate at steady-state (c.f. Eqns. 19 and 20) to obtain the community matrix, i.e., the Jacobian matrix evaluated at $(M^*, N^*)$. The equilibrium point $(M_i^*, N_j^*)$ is locally stable if and only if the dominant eigenvalue, $\lambda_{\max}$, the eigenvalue with the largest real part, has negative real part, i.e., $\mathrm{re}(\lambda_{\max}) < 0$.

##### Community matrix structure

The community matrix has a block structure:

$$J_\alpha\big|_{M^*,N^*} = \begin{bmatrix} \dfrac{\partial}{\partial M_i}\dfrac{dM_i}{dt}\Big|_{M^*,N^*} & \dfrac{\partial}{\partial N_j}\dfrac{dM_i}{dt}\Big|_{M^*,N^*} \\ \dfrac{\partial}{\partial M_i}\dfrac{dN_j}{dt}\Big|_{M^*,N^*} & \dfrac{\partial}{\partial N_j}\dfrac{dN_j}{dt}\Big|_{M^*,N^*} \end{bmatrix} \tag{22}$$

The diagonal elements of the community matrix, represent intraspecific effects:

$$\frac{\partial}{\partial M_i}\frac{dM_i}{dt}\Big|_{M_i^*,N_j^*} = \frac{\phi_{ij}\beta_i M_i^*}{(e_i + M_i^*)^2}\sum_{j\in S_A} N_j^* - d_i M_i^*, \tag{23}$$

$$\frac{\partial}{\partial N_j}\frac{dN_j}{dt}\Big|_{M_i^*,N_j^*} = -d_j N_j^*. \tag{24}$$

The off-diagonal elements capture interspecific interactions:

$$\frac{\partial}{\partial N_j}\frac{dM_i}{dt}\bigg|_{M_i^*,N_j^*} = \phi_{ij}\alpha_{ij}\frac{h_j M_i^*}{(h_j+N_j^*)^2} - \frac{\phi_{ij}\beta_i M_i^*}{e_i+M_i^*} \tag{25}$$

$$\frac{\partial}{\partial M_i}\frac{dN_j}{dt}\bigg|_{M_i^*,N_j^*} = \phi_{ji}\alpha_{ji}\frac{h_i N_j^*}{(h_i+M_i^*)^2} \tag{26}$$

In our case, the sign structure of the community matrix reflects the mutualistic nature of plant-pollinator interactions in a consumer-resource setting:

$$\text{sign}(J_\alpha\big|_{M^*,N^*}) = \begin{bmatrix} +/- & +/- \\ + & - \end{bmatrix} \tag{27}$$

Starting with interspecific effects, the bottom-left block contains only positive entries because pollinators only benefit from plants. By contrast, the top-right block can contain both positive and negative entries because plants experience both benefits (pollination services) and costs (resource allocation to rewards) from interactions. With intraspecific effects, the bottom-right block contains only negative entries, representing pollinator self-regulation. The situation is more complex with plants, with the top-left block in principle containing both positive and negative entries. This is because this block describes the effect of an (infinitesimally small) increase in focal plant density on focal plant dynamics and so, in addition to plant self-regulation, also includes the effects of changes in per capita resource production cost. Despite (infinitesimally) more individuals of the focal plant, the cost incurred for producing resources remains the same because the density of pollinators is not considered to change. As such, following the increase in focal plant density, we can consider there to be an effective reduction in cost, i.e., a reduction in negative effect, per capita, on the focal plant species, which can be understood as a positive contribution to the change in focal plant species abundance.

##### Distributional analysis

We compute the dominant eigenvalue $\lambda_{\max}$ for each of the 100,000 sampled interaction coefficient sets. The resulting distribution of eigenvalues describes the stability of the system across a wide range of feasible sets of interaction coefficients.

A smooth density concentrated to the left of the imaginary axis (i.e., $\text{re}(\lambda_{\max}) < 0$) indicates robust stability and adequate sampling. An irregular density distribution suggests insufficient sampling, requiring additional samples. If a substantial portion of the density crosses the imaginary axis (i.e., $\text{re}(\lambda) \geq 0$), we adjust the tuning parameters $\varepsilon_f$ and $\varepsilon_s$ to shift the death rates $d_i$ and $d_j$ to increase system stability.

### H: Sensitivity to phenology

While the community matrix characterizes how growth rates and therefore species densities respond to perturbations in population densities at the steady state, it does not capture responses to changes in interaction timing, i.e., phenology. The phenological overlap coefficients $\phi_{ij}$ provide a mechanism to assess how shifts in temporal overlap affect population dynamics.

#### The phenological sensitivity matrix

We introduce a matrix $B$ that captures species responses to changes in phenological overlap, analogous to how the community matrix above captures responses to direct perturbations to species densities at the steady state. In contrast, each entry of $B$ quantifies how much a species' growth rate changes at the steady state when a particular interaction becomes less temporally overlapping.

Formally, let $\{\phi\}$ denote the set of all non-zero phenological overlap coefficients $\phi_{ij}$ for interacting species pairs $(i,j)$ where $i \in S_P$ and $j \in S_A$. The matrix $B$ has dimensions $|S_P| + |S_A|$ by $|\{\phi\}|$, with one row for each species and one column for each phenological overlap coefficient. Each entry is defined as

$$B_{lk} = -\frac{\partial}{\partial \phi_k}\left(\frac{dX_l}{dt}\right)\bigg|_{M_i^*,N_j^*} \tag{28}$$

where $X_l = M_l$ if $l \in S_P$ and $X_l = N_l$ if $l \in S_A$, $\phi_k$ is the $k$-th element of $\{\phi\}$, and the partial derivative is evaluated at steady-state densities. The negative sign reflects our focus on decreases in phenological overlap.

For plant species $i$ interacting with pollinator $j$, the entries are

$$B_{ik} = -\frac{\partial}{\partial \phi_{ij}} \left( \frac{dM_i}{dt} \right) \Bigg|_{M_i^*, N_j^*} = - \left[ \alpha_{ij} \frac{M_i^* N_j^*}{h_j + N_j^*} - \beta_i \frac{M_i^* N_j^*}{e_i + M_i^*} \right] \tag{29}$$

For pollinator species $j$ interacting with plant $i$, the entries are

$$B_{jk} = -\frac{\partial}{\partial \phi_{ji}} \left( \frac{dN_j}{dt} \right) \Bigg|_{M_i^*, N_j^*} = -\alpha_{ij} \frac{M_i^* N_j^*}{h_i + M_i^*} \tag{30}$$

These derivatives are computed using the sampled interaction coefficients $\alpha_{ij}$ and observed steady-state densities.

#### Differences between the community matrix and the $B$ matrix

There are three major differences between the community matrix, Eqns. 23–26, and the $B$ matrix, Eqns. 29 and 30. First, by design, the two types of matrix measure the effects of two different types of change: species densities—amount—in the case of the community matrix and interaction timing—temporal availability—in the case of the $B$ matrix. Second, the community matrix includes intraspecific effects (e.g., self-regulation) as well as interspecifc effects, whereas the $B$ matrix is focused on interspecific effects and as such its entries do not include species-specific death (and growth) rate parameters. Third, entries scale with species densities differently, with entries in the $B$ matrix scaling with the "mass action" product of steady-state densities, $M_i^* N_j^*$, consistent with the $B$ matrix's focus on the effects on changes in interaction availability rather than changes in species densities. Taken together, although the entries of the two types of Jacobian matrices evaluated at steady state include similar parameters, there are sufficient formal differences between the community matrix and $B$ matrix that when dominant eigenvalues (from community matrices) and dominant singular values (from $B$ matrices) are computed for the same system and set of parameters, there should be no mathematically-based expectation that the two derived quantities will be strongly correlated with one another.

#### Measuring phenological sensitivity

Unlike the square community matrix, the $B$ matrix is generally rectangular (more phenological overlap coefficients than species). Therefore, we cannot use eigenvalues to characterize its properties. Instead, we employ the largest singular value $\sigma_{\max}(B)$ as a measure of phenological sensitivity.

The largest singular value quantifies the maximum amplification of perturbations: if phenological overlaps change by a small amount, $\sigma_{\max}(B)$ bounds how much species growth rates can change. Larger singular values indicate greater (maximum) sensitivity to phenological changes. Unlike the dominant eigenvalue of the community matrix, singular values are always non-negative and do not correspond to stability, instead they measure the magnitude of the response. The relationship between dominant singular values of the $B$ matrix and dominant eigenvalues of the community matrix for the three field sites are shown in Figs. S1–S3.

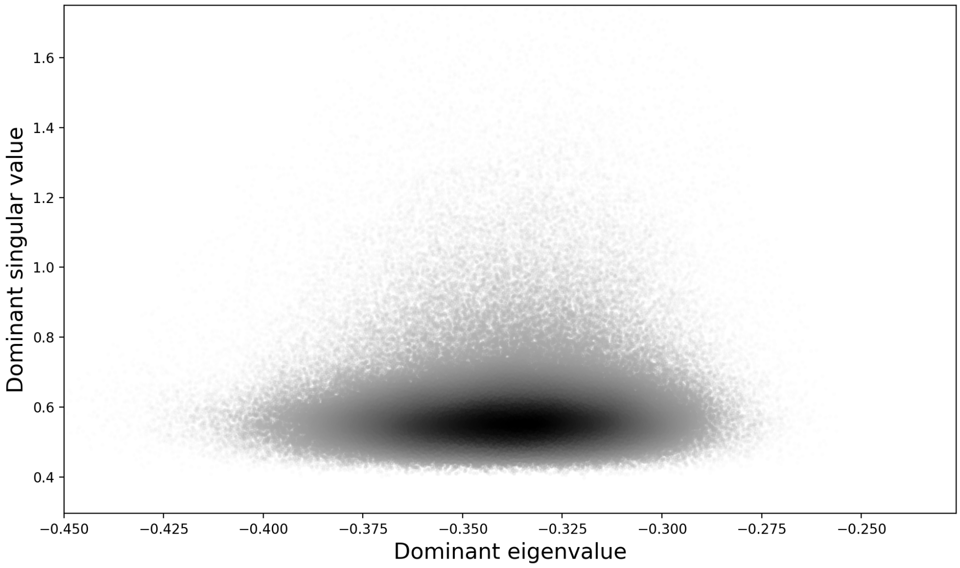

**Figure S1:** Relationship between dominant singular values of the $B$ matrix and dominant eigenvalues of the community matrix for the Avery Picnic ("AP") site (Spearman's rank correlation: $r = 0.051$). Each point in the scatter plot represents a dominant singular value and dominant eigenvalue pair corresponding to a distinct set of sampled interaction strengths ($n = 100,000$ samples); darker areas indicate greater densities of points.

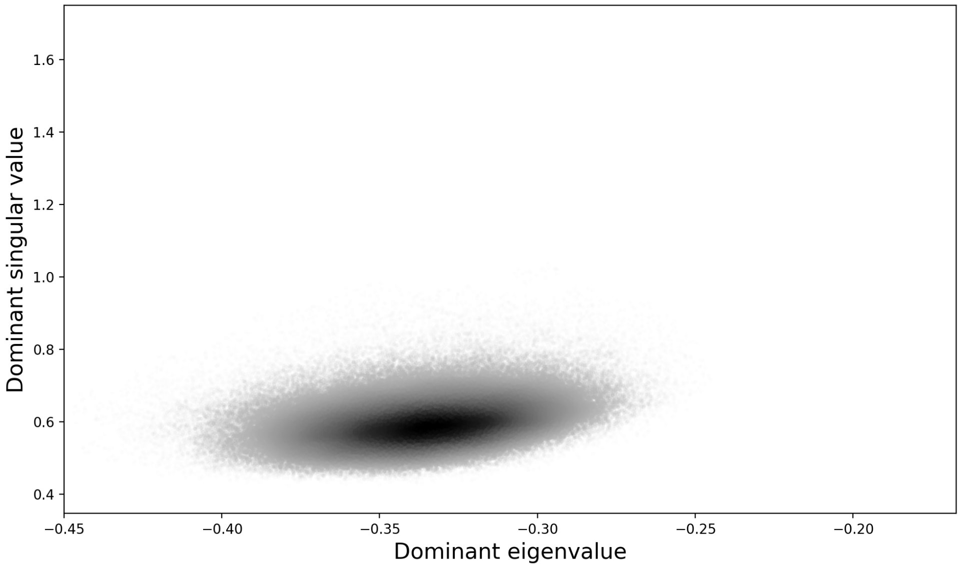

**Figure S2:** Relationship between dominant singular values of the $B$ matrix and dominant eigenvalues of the community matrix for the Gothic Town ("GT") site (Spearman's rank correlation: $r = 0.297$). Each point in the scatter plot represents a dominant singular value and dominant eigenvalue pair corresponding to a distinct set of sampled interaction strengths ($n = 100,000$ samples); darker areas indicate greater densities of points.

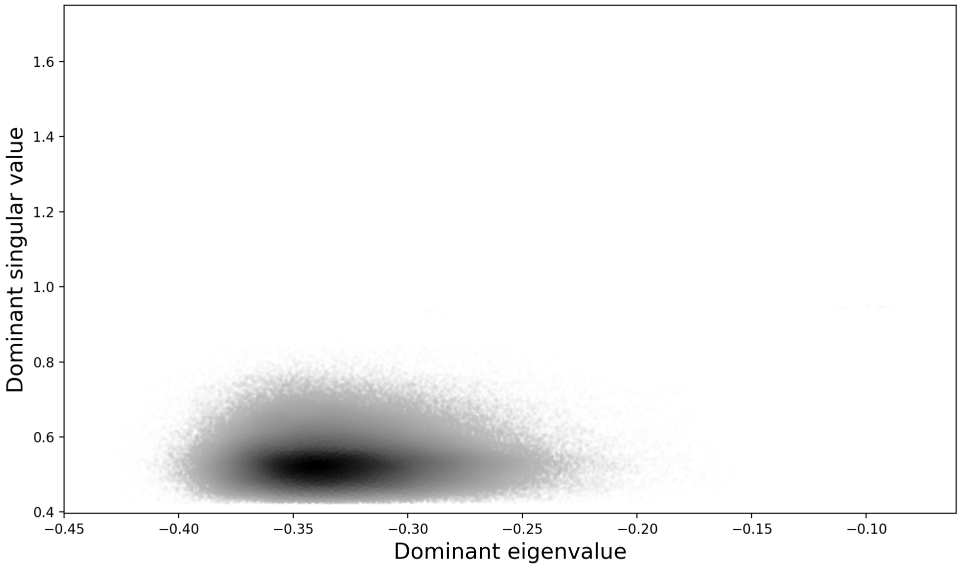

**Figure S3:** Relationship between dominant singular values of the $B$ matrix and dominant eigenvalues of the community matrix for the Virginia Basin ("VB") site (Spearman's rank correlation: $r = 0.008$). Each point in the scatter plot represents a dominant singular value and dominant eigenvalue pair corresponding to a distinct set of sampled interaction strengths ($n = 100,000$ samples); darker areas indicate greater densities of points.

#### Visualization

To visualize phenological sensitivities at the level of individual interactions, we reorganized the $B$ matrix into two matrices that preserve the underlying bipartite network structure. Since the effect of an interaction is asymmetric ($\phi_{ij} \neq \phi_{ji}$), we constructed separate matrices from the perspective of plant and pollinator species. For each of the 100,000 sampled sets of interaction coefficients, we computed the per capita phenological sensitivity matrix by dividing each entry of the $B$ matrix by the corresponding species density, then averaged these matrices across all samples. Per capita sensitivities represent the proportional change in a species' growth rate per unit decrease in phenological overlap, enabling comparisons among species with different baseline population sizes. The rearranged $B$ matrices for the three field sites are shown in Figs. S4–S6.

**Matrix construction.** For each interaction between plant $i$ and pollinator $j$, we created a plant-perspective matrix $B_P$ of dimensions $|S_P| \times |S_A|$, where each entry $(i, j)$ contains the per capita sensitivity value corresponding to $\phi_{ij}$ (the effect on plant $i$ when overlap with pollinator $j$ decreases). We also created a pollinator-perspective matrix $B_A$ of the same dimensions, where each entry $(i, j)$ contains the per capita sensitivity value corresponding to $\phi_{ji}$ (the effect on pollinator $j$ when overlap with plant $i$ decreases). Non-interacting species pairs have entries equal to zero in both matrices.

**Interpreting the sensitivity matrices.** These matrices reveal multiple aspects of phenological vulnerability. First, interaction-level sensitivity: each element $(i, j)$ quantifies how vulnerable the focal species is to phenological changes in the particular interaction between plant $i$ and pollinator $j$, with large values indicating interactions that are critical for species persistence. Second, species-level sensitivity: row sums in $B_P$ (for plants) and column sums in $B_A$ (for pollinators) quantify the total sensitivity of each species to phenological changes across all its interaction partners, with species that have large sums being especially vulnerable to phenological changes. Third, mean sensitivity: the average sensitivity across all interactions for each species provides a standardized measure of vulnerability that accounts for differences in the number of interaction partners.

In our visualizations, we display these two matrices side-by-side: the left panels show sensitivities from the plant perspective ($B_P$), and the right panels show sensitivities from the pollinator perspective ($B_A$). An additional column in each panel summarizes the mean per capita sensitivity of each species, averaged over all its interactions. These species-level per capita mean sensitivities were used as the response variable in the regression analysis.

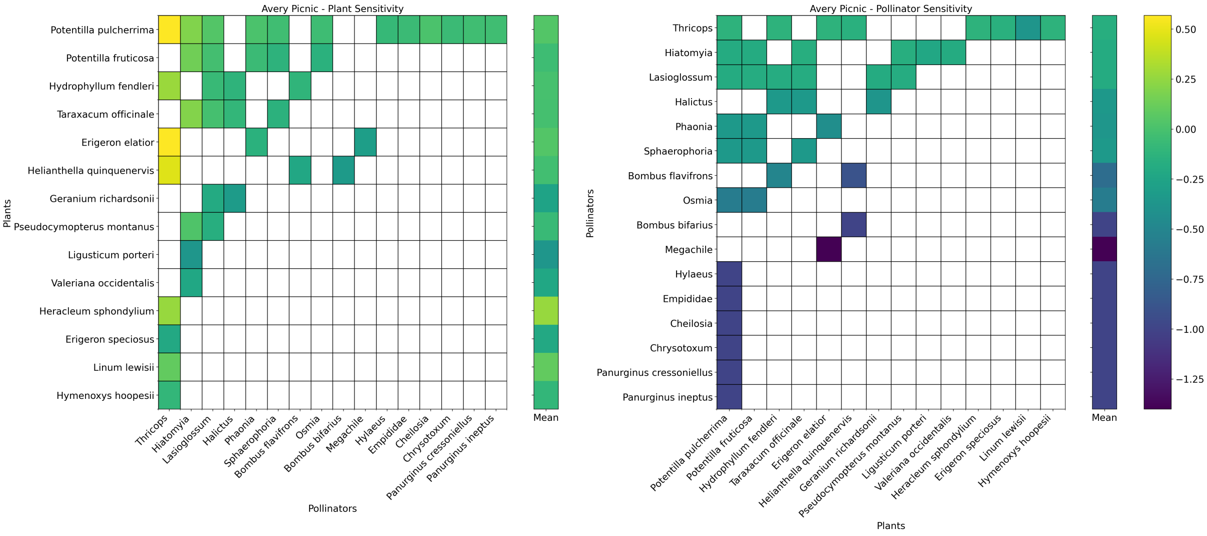

**Figure S4:** Rearranged $B$ matrix for the Avery Picnic ("AP") site.

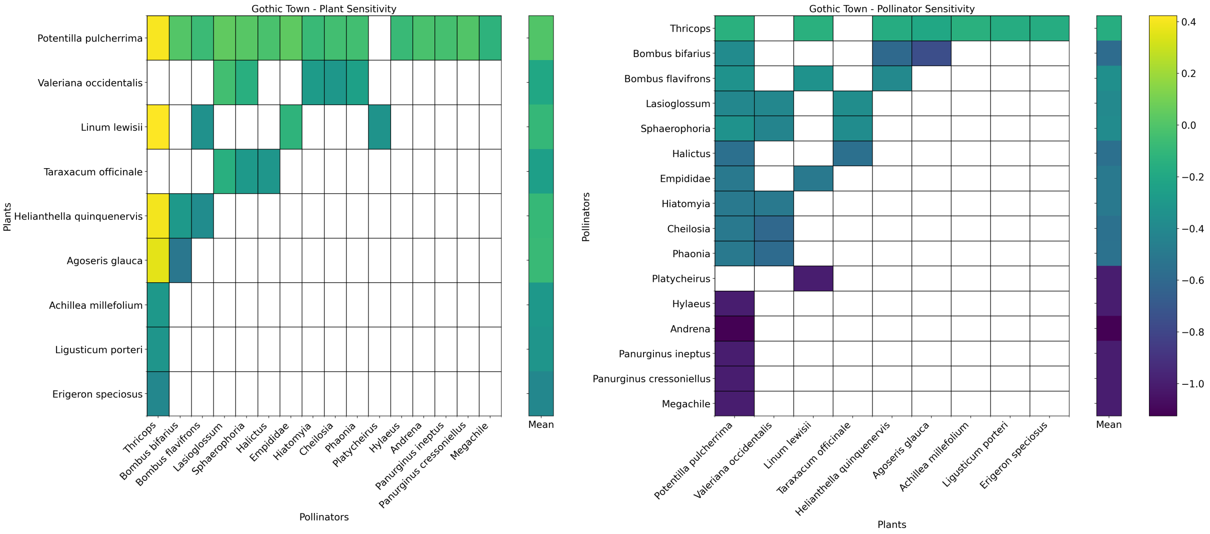

**Figure S5:** Rearranged $B$ matrix for the Gothic Town ("GT") site.

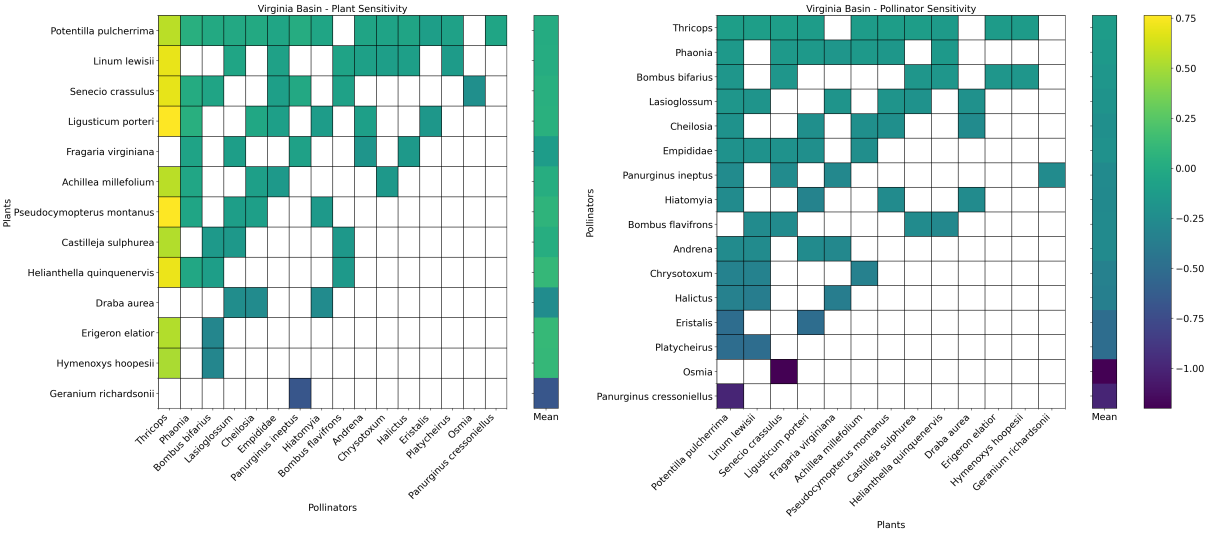

**Figure S6:** Rearranged $B$ matrix for the Virginia Basin ("VB") site.

### References

[1] C. Morozumi, X. Loy, V. Reynolds, A. Schiffer, B. Morrison, J. Savage, and B. J. Brosi, "Simultaneous niche expansion and contraction in plant-pollinator networks under drought," *Oikos*, vol. e09265, 2022.

[2] J. Holland and D. DeAngelis, "A consumer-resource approach to the density-dependent population dynamics of mutualism," *Ecology*, vol. 91, pp. 1286–1295, 2010.

[3] S. Allesina and S. Tang, "The stability-complexity relationship at age 40: A random matrix perspective," *Popul. Ecol.*, vol. 57, pp. 63–75, 2015.

[4] D. Kaufman and R. Smith, "Direction choice for accelerated convergence in hit-and-run sampling," *Oper. Res.*, vol. 46, pp. 84–95, 1998.

[5] P. J. Goulart and Y. Chen, "Clarabel: An interior-point solver for conic programs with quadratic objectives; eprint arxiv:2405.12762," 2024.
